# Direct type I Interferon signaling in hepatocytes control malaria

**DOI:** 10.1101/2022.01.26.477934

**Authors:** Camila Marques-da-Silva, Kristen Peissig, Michael P. Walker, Justine Shiau, Dennis E. Kyle, Rahul Vijay, Scott E. Lindner, Samarchith P. Kurup

## Abstract

Malaria is a devastating disease impacting over half of the world’s population. *Plasmodium* parasites that cause malaria undergo obligatory development and replication in hepatocytes before infecting red blood cells and initiating clinical disease. While type I interferons (IFNs) are known to facilitate innate immune control to *Plasmodium* in the liver, how they do so has remained unresolved, preventing the manipulation of such responses to combat malaria. Utilizing transcriptomics, infection studies, and a novel transgenic *Plasmodium* strain that exports and traffics Cre recombinase, we show that direct type I IFN signaling in *Plasmodium*-infected hepatocytes is necessary and sufficient to control malaria. We also show that the majority of infected hepatocytes naturally eliminate *Plasmodium* infection, revealing the potential existence of anti-malarial cell-autonomous immune responses in such hepatocytes. These discoveries challenge the existing paradigms in *Plasmodium* immunobiology and are expected to inspire new anti-malarial drugs and vaccine strategies.

## Introduction

Clinical malaria infections caused by *Plasmodium* parasites are responsible for the death and suffering of millions of people around the globe (WHO, 2019). The sporozoite stage of *Plasmodium* inoculated into humans by *Anopheles* mosquitoes has to undergo developmental transformation and replication in hepatocytes (constituting the ‘liver stage’ of malaria), before being able to infect red blood cells and causing the highly symptomatic and dangerous ‘blood stage’ of malaria (Cowman et al., 2016). The blood stage is exclusively responsible for all of the morbidity and mortality associated with malaria, as well as its transmission. Therefore, limiting the progression of *Plasmodium* beyond the liver is the primary goal of the majority of current strategies aimed at controlling or eradicating malaria (Mo and McGugan, 2018; Mo et al., 2020). However, a major impediment to such approaches is how little we know about the nature of host responses to malaria infection in the liver (Gazzinelli et al., 2014). Type I IFNs are the only known mediators of innate immune control of liver-stage malaria (Gazzinelli *et al*., 2014; Liehl et al., 2014; Miller et al., 2014). There is increasing evidence that limiting *Plasmodium* infection in the liver by induction of type I IFNs blunts the extent and impact of clinical malaria (He et al., 2020). However, how type I IFNs drive the control of *Plasmodium* in the liver has remained unresolved (Liehl *et al*., 2014; Miller *et al*., 2014). Currently, it is presumed that type I IFNs produced by *Plasmodium*-infected hepatocytes facilitate the recruitment of immune cells from circulation either through direct signaling or through the induction of secondary chemokine signals via ‘bystander’ uninfected hepatocytes, and that these immune cells would eliminate the *Plasmodium*-infected hepatocytes through some unknown mechanism (Gazzinelli *et al*., 2014; Liehl *et al*., 2014). In contrast to these notions, using a novel Cre recombinase expressing *Plasmodium* strain capable of selectively ablating type I IFN signaling exclusively in the infected hepatocytes, we show that type I IFNs directly signal the infected hepatocytes to bring about control of malaria infection in the liver. This finding should encourage additional research into IFN-induced cell-autonomous immune mechanisms targeting *Plasmodium* in hepatocytes, and stimulate novel therapeutic opportunities and vaccination strategies against malaria.

## Results

### Type I Interferon signaling evident in *Plasmodium*-infected hepatocytes

It is well known that type I IFNs are critical for controlling malaria infection in the liver, and systemic induction of adequate type I IFNs significantly limits *Plasmodium* development in the liver (Figures 1A-B, S1A) (Gazzinelli *et al*., 2014; Liehl *et al*., 2014; Miller *et al*., 2014). A direct role for type I IFNs in targeting *Plasmodium* or *Plasmodium*-infected hepatocytes had been previously ruled out, based on the observation that treatment of *P. berghei*-infected primary hepatocytes *in vitro* with the drug DMXAA, a potent inducer of type I IFNs, did not impact *Plasmodium* development (Liehl *et al*., 2014). However, the primary pathway through which DMXAA treatment induces type I IFNs in mouse cells is through the stimulation of the cytosolic receptor, STING (Stimulator of interferon genes) (Conlon et al., 2013). It was later revealed that STING is not expressed in hepatocytes (Thomsen et al., 2016). Therefore, DMXAA treatment would not have induced type I IFNs in primary hepatocyte cultures. Of note, *Plasmodium* nucleic acids, which can potentially stimulate STING are known to gain access to the cytosol of infected hepatocytes (Ni et al., 2018; Slavik et al., 2021). Yet, genetic deficiency of STING nor its signaling partner cGAS did not alter *Plasmodium* control in the liver, further supporting the absence of STING signaling in hepatocytes or its direct relevance to liver-stage malaria (Figure S1B). In contrast, direct IFNα and IFNβ (IFNα/β) treatment of *P. falciparum* infected human primary hepatocytes or *P. berghei* infected murine primary hepatocytes induced a significant, dose-dependent control of the infection (Figure 1C-D). These findings renewed our interest in the possibility that type I IFNs directly mediated control of *Plasmodium* in hepatocytes.

**Figure 1:**
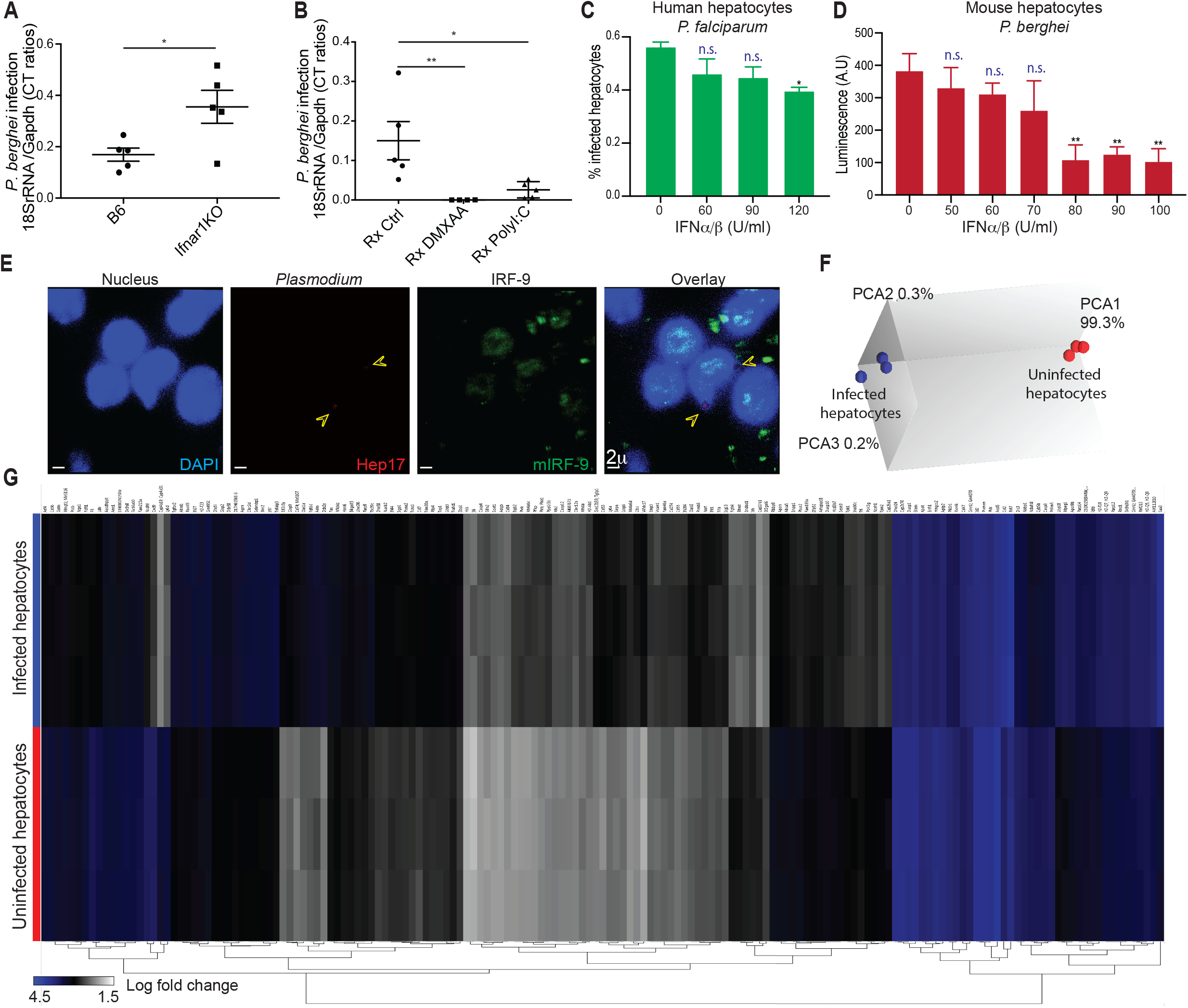
Type I interferon signaling in *Plasmodium*-infected hepatocytes. (A), Scatter plots showing relative liver-parasite burdens in the indicated mice inoculated with 3×10^4^*Pb* spz, 36h p.i. (B), Scatter plots showing relative liver-parasite burdens at 36h p.i. in B6 mice treated with vehicle control, STING agonist DMXAA, or TLR3 agonist polyI:C (−1d p.i.), and inoculated with 3×10^4^*Pb* spz. A-B, Dots represent individual mice, data presented as mean + s.e.m and analyzed using 2-tailed t-tests (A) or ANOVA with Tukey’s correction (B). (C) Frequency of *P. falciparum* infected primary human hepatocytes treated with the indicated concentrations of IFNα/β, 4d p.i. (D) Parasite loads indicated by luciferase activity in primary mouse hepatocytes infected with *Pb-luc* and treated with the indicated concentrations of IFNα/β, 36h p.i. C-D Data presented as mean + s.e.m and analyzed using ANOVA comparing the indicated groups with the group treated with 0U/ml IFNα/β. (E) Representative pseudo-colored confocal images depicting IRF-9 translocation into host cell nuclei, indicating type-I IFN signaling in *Pb* infected B6 mouse liver, at 24h p.i. The arrows indicate *bona fide* liver-stage *Pb* stained for Hep17 protein in its parasitophorous vacuolar membrane (PVM) in uninfected or *Pb* infected primary mouse hepatocytes. (F) Principal component analysis representing gross transcriptional differences between *Pb* infected or naive B6 hepatocytes *in vitro*. (G) Transcriptional perturbations in interferon regulated genes alone in *Pb* infected or naive B6 hepatocytes *in vitro*. A-E represent ≥3 separate experimentsbiological replicates. * P<0.05, **P<0.01, n.s P≥0.05

Considering that a variety of IFN-induced genes and pathways are known to direct host cell elimination of intracellular pathogens (McNab et al., 2015), we hypothesized that autocrine or paracrine type I IFN signaling in hepatocytes would drive clearance of *Plasmodium* from hepatocytes. In support of this premise, we observed IRF9 translocation into the nuclei of *P. berghei*-infected hepatocytes and other neighboring cells in the livers of infected mice (Figure 1E), indicating active type I IFN signaling in these cells (MacMicking, 2012). Microarray analysis in *P. berghei*-infected primary mouse hepatocytes indicated that a large number of type I IFN responsive genes were transcriptionally altered in *Plasmodium*-infected hepatocytes (Figure 1F-G). Together, these data demonstrate that type I IFN signaling is active in *Plasmodium*-infected hepatocytes.

### Type I Interferon signaling in infected hepatocytes controls malaria infection

To determine the extent to which type I IFN signaling in infected hepatocytes contributed to the overall control of liver-stage malaria, we sought to negate type I IFN signaling solely in the infected hepatocytes. To this end, we generated a strain of *P. berghei, Pb-Cre*, which expresses and exports Cre recombinase into infected host hepatocytes wherein it may excise DNA sequences flanked by loxP (locus of X-over P1) sites (‘floxed’). *Pb-Cre* parasites were generated by fusing Cre recombinase with an SV40 nuclear localization signal (NLS) and HA tag on the C-terminus of Lisp2 by CRISPR-mediated insertion of these coding sequences in the endogenous *pblisp2* genomic locus (Figure S2A). Lisp2 protein is expressed by *bona fide* liver stages (merozoites) of *Plasmodium* in hepatocytes and is known to be transported across the parasitophorous vacuolar membrane into the hepatocyte cytosol (Gupta et al., 2019; Orito et al., 2013; Woodard et al., 2010). Therefore, the fusion with Lisp2 would limit Cre expression to merozoites and facilitate its transport into the hepatocyte cytosol, where the SV40 NLS would drive its import into the host hepatocyte nucleus (Figure S2B). To demonstrate the extent to which *Pb-Cre* can ablate target DNA sequences in host hepatocytes, we infected primary hepatocytes derived from Ai14 mice with *Pb-Cre*. Ai14 mice possess a floxed STOP cassette preventing the transcription of a CAG promoter-driven red fluorescent protein variant, tdTomato (Madisen et al., 2010). Upon excision of the STOP cassette, tdTomato expression is ‘turned on’ in Ai14 cells. Ai14 hepatocytes infected with *Pb-Cre* showed robust tdTomato expression (Figure S2C), indicating that *Pb-Cre* can be reliably used to ablate floxed target DNA sequences in *Plasmodium*-infected cells. Therefore, *Pb-Cre* provided an opportunity to determine the relevance of specific genes in host hepatocytes in the biology of *Plasmodium* infection.

To investigate how type I IFN responses specifically in the *Plasmodium*-infected hepatocytes impacted the overall control of liver-stage malaria, we utilized Ifnar1^fl^ mice, in which the exon 3 of type I interferon α/β receptor (IFNAR) is flanked by loxP sequences (Kamphuis et al., 2006). As expected, *Pb-Cre* infection induced tangible loss of IFNAR expression in the Ifnar1^fl^ hepatocytes (Figure 2A). Of note, the time window available to observe a complete loss of gene expression in hepatocytes is limited by the ~48-hour lifespan of *Plasmodium* development in mouse hepatocytes. Nevertheless, even partial reduction in gene expression in parenchymal cells achieved using tools such as siRNA has provided ground-breaking advancements in our understanding of host gene functions in a variety of liver diseases, including in malaria (Canal et al., 2015; Liehl *et al*., 2014; Rudalska et al., 2014). Compared to the control B6 hepatocytes, Ifnar1^fl^ hepatocytes infected with *Pb-Cre* failed to optimally control the infection *in vitro* (Figures 2B, S2D). This outcome was also noticed *in vivo*, where, in comparison to control B6 mice, Ifnar1^fl^ mice failed to limit *Pb-Cre* infection in the liver even with systemic induction of type I IFNs via DMXAA administration (Figure 2C). Systemic delivery of DMXAA in mice is known to induce type I IFNs through the stimulation of STING in a variety of cell-types such as the dendritic cells and macrophages (Curran et al., 2016). Sufficient administration of exogenous IFNα/β during the liver stage also significantly limited the progression *Pb-Cre* to blood stage infection in B6 mice, but not in IFNAR^fl^ mice (Figure 2D), demonstrating the relevance of IFN signaling in hepatocytes in determining clinical outcomes. These findings showed that type I IFN signaling specifically and directly in *Plasmodium*-infected hepatocytes was instrumental in limiting malaria infection. Consistent with this notion, the liver parasite burdens in *Pb-Cre* infected Ifnar1^fl^ mice were similar to those in control *Pb-Ova* (*P. berghei* transgenically expressing Ovalbumin) infected Ifnar1^fl^-AlbCre mice (Figure 2C), which lack IFNAR in all hepatocytes (Liehl *et al*., 2014). This finding implied that the contribution of type I IFN signaling in uninfected hepatocytes to the control of malaria infection in the liver is negligible. Taken together, these data indicated that type I IFN signaling in *Plasmodium*-infected hepatocytes is both necessary and sufficient to bring about IFN mediated control of malaria infection in the liver.

**Figure 2:**
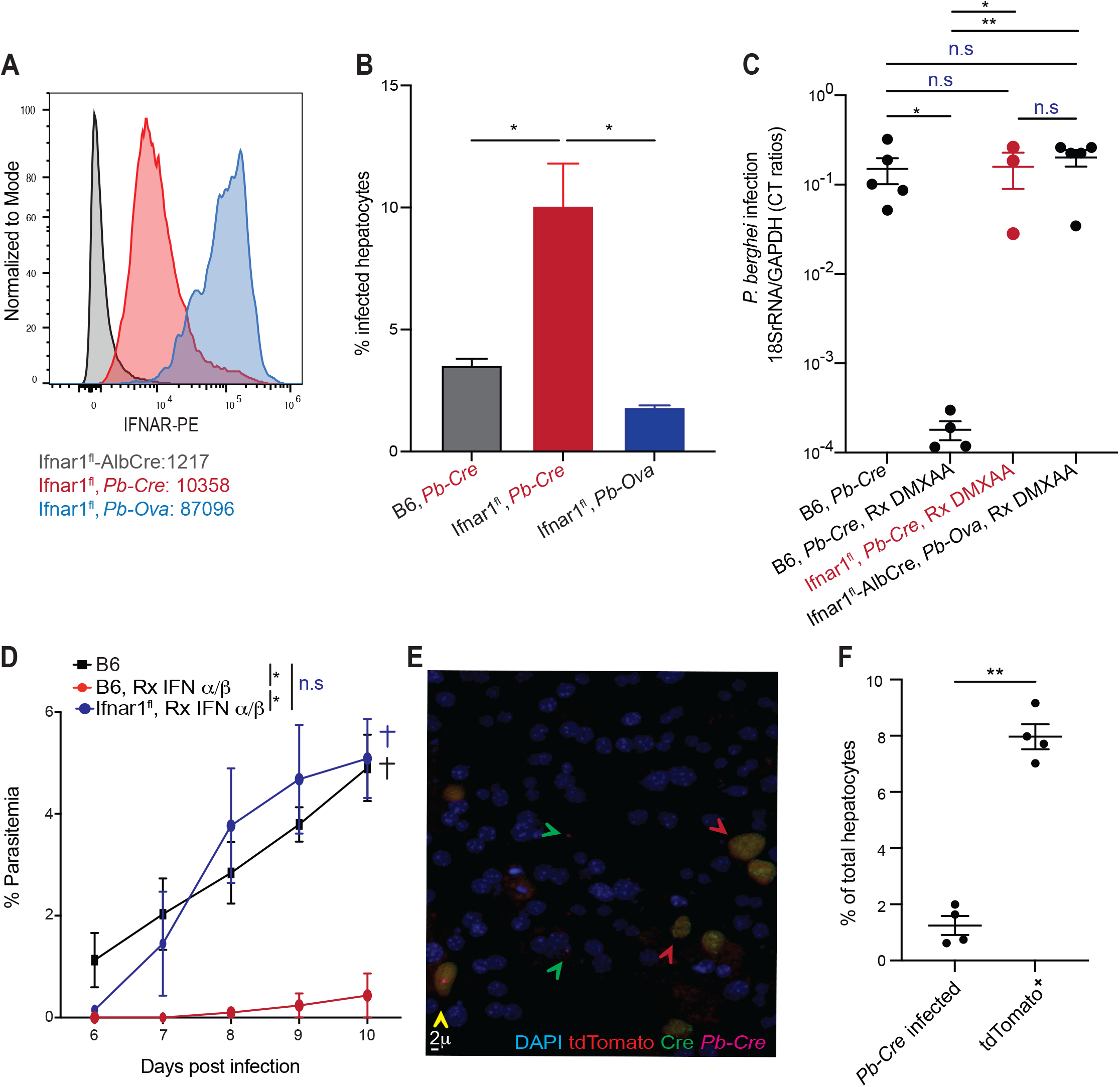
Type I IFN signaling in the infected hepatocytes control malaria infection. (A) Histograms representing the expression levels of IFNAR in *Pb-Cre* infected Ifnar1^flox^ hepatocytes. *Pb-Ova* infected Ifnar1^flox^ hepatocytes or uninfected Ifnar1^flox^-AlbCre hepatocytes served as controls. CellTrace Violet used to stain sporozoites to identify the infected hepatocytes. (B) Frequencies of infected B6 or Ifnar1^flox^ primary hepatocytes infected with *Pb-Cre* or control *Pb-Ova* parasites when co-incubated for 36h. (C) Scatter plots showing relative liver-parasite burdens at 36h p.i. in the indicated groups of mice inoculated with 3×10^4^ *Pb-cre* or *Pb-ova* sporozoites, and treated with or without DMXAA as indicated (1d p.i.). Dots represent individual mice. (D) Kinetics of parasitemia in the indicated mice inoculated with *Pb-Cre* sporozoites and treated with or without IFNα/β at 12h and 24h p.i. (E) Representative pseudo-colored confocal images indicating tdTomato expression, Cre localization or *Pb-Cre* in primary Ai14 mouse hepatocytes. Green arrows indicate *Pb-Cre* infected hepatocytes, red arrow indicates tdTomato expressing hepatocytes and yellow arrow shows tdTomato expressing cells that have detectable *Pb-Cre* (Hep17^+^) parasites in them. See Extended Data Fig. 2e for individual channels. (F) Scatter plot indicating the frequency of primary Ai14 mouse hepatocytes co-incubated with *Pb-Cre* sporozoites showing tdTomato expression or detectable *Pb-Cre* infection. Data presented as mean + s.e.m and analyzed using ANOVA with Tukey’s correction (B-D) or 2-tailed t-tests (F). All data represent ≥3 separate experiments. * P<0.05, ** P <0.01, n.s P≥0.05

The induction of tdTomato expression in Ai14 hepatocytes by *Pb-Cre* infection presented the unique opportunity to identify infected hepatocytes without relying on the detection of proteins expressed by the parasite. Given that Cre-based activation of tdTomato expression in infected Ai14 hepatocytes is permanent, this system also enabled the identification of previously infected hepatocytes that may have cleared *Pb-Cre* infection. Surprisingly, a large number of Ai14 hepatocytes co-incubated with *Pb-Cre in vitro* expressed tdTomato without the presence of detectable parasites in them (Figure 2E-F, S2E). Considering that only *bona fide* liver stages of *Pb-Cre* would express Cre (Gupta *et al*., 2019; Orito *et al*., 2013), and that baseline tdTomato expression in Ai14 hepatocytes is negligible (Figure S2F), our findings suggested that the majority of hepatocytes infected with *Pb-Cre* eliminate the infection. Furthermore, Cre recombinase remained detectable in the majority of tdTomato expressing hepatocytes, suggesting that *Pb-cre* were probably present in these at some point. The tdTomato expression in such cells is also unlikely to be spontaneous or background (Figure S2E). Some hepatocytes, which are possibly in their early stages of *Pb-Cre* infection are not expected to show detectable levels of Cre or tdTomato expression, considering that Lisp2 expression is limited to the mid-late merozoite stage of *Plasmodium* in hepatocytes (Gupta *et al*., 2019; Orito *et al*., 2013). Together, these findings implied the possibility of active cell-intrinsic immune mechanisms being present in hepatocytes, which aid in limiting *Plasmodium* infection in the liver.

## Discussion

We show that innate immune control of *Plasmodium* in its liver stage occurs through type I IFN signaling specifically in the infected hepatocytes, and that the majority of the hepatocytes eliminate *Plasmodium* parasites in the liver without tangible contributions from uninfected hepatocytes or other immune cells. This is contrary to the existing notion in the field that type I IFNs produced by *Plasmodium* infected hepatocytes would stimulate uninfected hepatocytes to induce chemokine signals that recruit leukocytes from circulation, which subsequently eliminate the infected hepatocytes (Gazzinelli *et al*., 2014; He *et al*., 2020; Liehl *et al*., 2014). Our findings indicate a direct type I IFN mediated elimination of *Plasmodium* in hepatocytes, potentially through autocrine and/ or paracrine signaling. We also show that such control of *Plasmodium* infection in the liver can have a significant impact on the onset and the overall parasite burden in the subsequent blood stage of malaria, indicating the relevance of type I IFN signaling in the infected hepatocytes to clinical outcomes.

Our data also suggest that cell-intrinsic immune responses in hepatocytes may be instrumental in routinely eliminating the majority of *Plasmodium* in infected hepatocytes. Although type I IFNs are considered key drivers of immune responses induced by or in immune cells, the roles of IFN-induced pathways in controlling infections in ‘non-immune’ cell-types are increasingly being appreciated (Gaudet et al., 2021; MacMicking, 2012). Given that type I IFNs are the only known drivers of innate immune control of *Plasmodium* in the liver, our work offers important insights into how malaria infection is naturally limited in the liver. There is emerging evidence for type I IFN responses impeding anti-malarial vaccine efficacies, as well as facilitating the elimination of dormant *P. vivax* hypnozoites in the liver capable of seeding relapsing malaria infections (Mancio-Silva et al., 2021; Minkah et al., 2019). Understanding how type I IFNs drive the elimination of *Plasmodium* from the infected hepatocytes might provide key insights into the mechanisms enabling these. Our findings provide a platform to investigate type I IFN signaling, as well as the cell-autonomous immune mechanisms it stimulates in hepatocytes that bring about natural control of *Plasmodium* infection in the liver.

### Limitations of the study

There is a possibility that *Pb-Cre* infection may not have induced uniform deletion of the IFNAR1 gene or the complete loss of IFNAR protein expression in all infected hepatocytes. However, reducing protein expression has often produced tangible phenotypic outcomes, providing valuable information about the functional roles of targeted genes in hepatocytes. (Canal *et al*., 2015; Liehl *et al*., 2014; Rudalska *et al*., 2014). Therefore, we anticipate *Pb-Cre* to serve as an important tool to dissect the contributions of any hepatocytes gene towards anti-malarial immunity going forward.

Another potential limitation of our study is the prospect of tdTomato expression being ‘turned on’ in Ai14 mouse hepatocytes without infection with *Pb-Cre*, due to Cre somehow gaining access to the nucleus of such hepatocytes. While hepatocytes are known to phagocytose exogenous particles (Soji et al., 1992), we do not anticipate extracellular Cre gaining access to hepatocyte nuclei following phagocytic uptake by hepatocytes. Furthermore, in all instances of tdTomato expression being observed in Ai14 mouse hepatocytes, we detected diffused Cre-recombinase in the hepatocyte cytoplasm, suggesting that these cells were likely previously infected by *Pb-Cre*.

---

**Figure S1:**
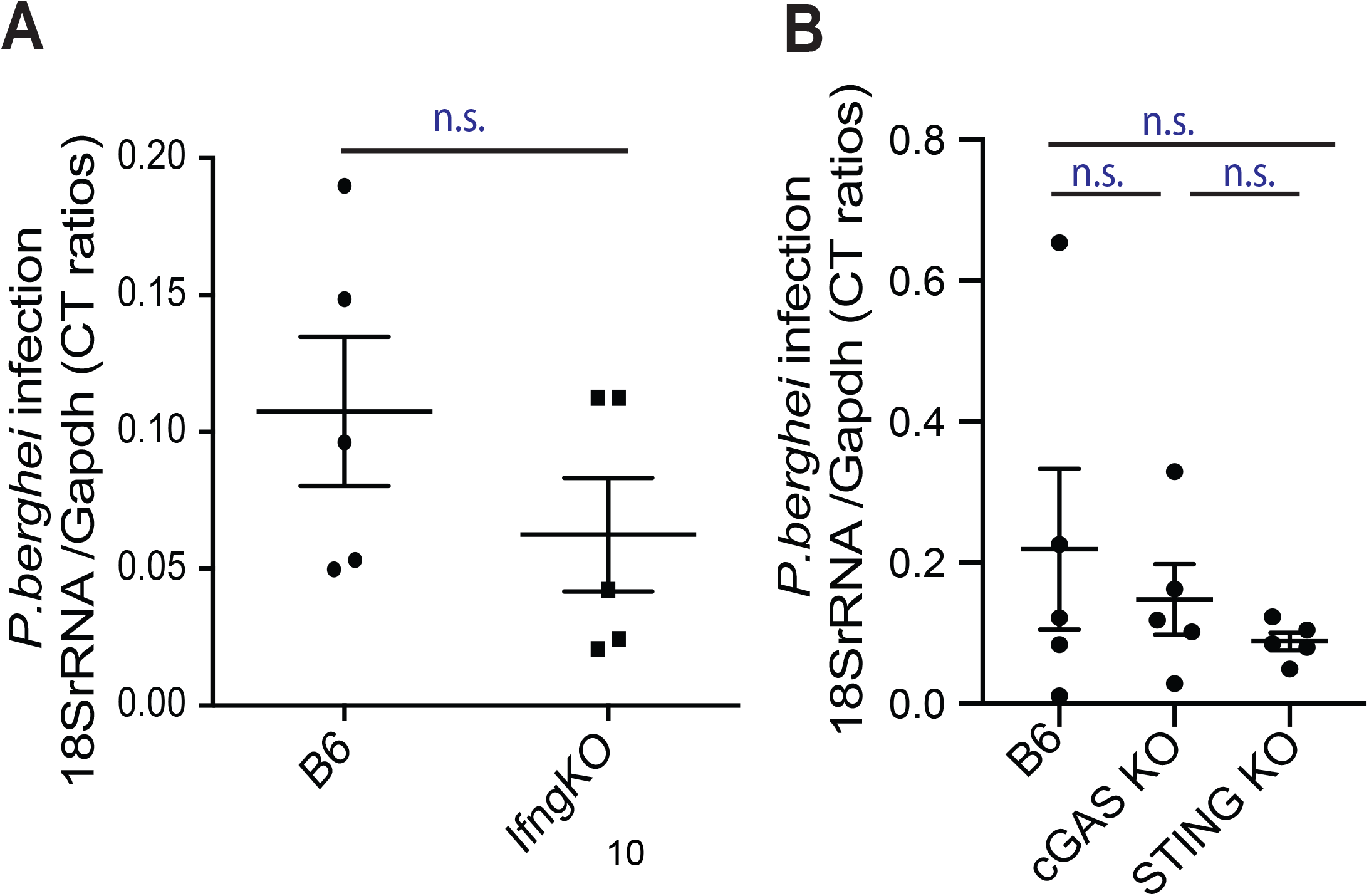
IFNγ or cGAS/STING do not impact control of *Plasmodium* infection in liver. (A-B) Scatter plots showing relative liver-parasite burdens in B6, or mice deficient in IFNγ (A), cGAS or STING (B), when inoculated with 3×10^4^*Pb* spz, determined at 36h p.i. Dots represent individual mice, data presented as mean + s.e.m and analyzed using 2-tailed t-tests (A) or ANOVA with Tukey’s correction (B). All data represent ≥3 separate experiments. n.s P≥0.05

**Figure S2:**
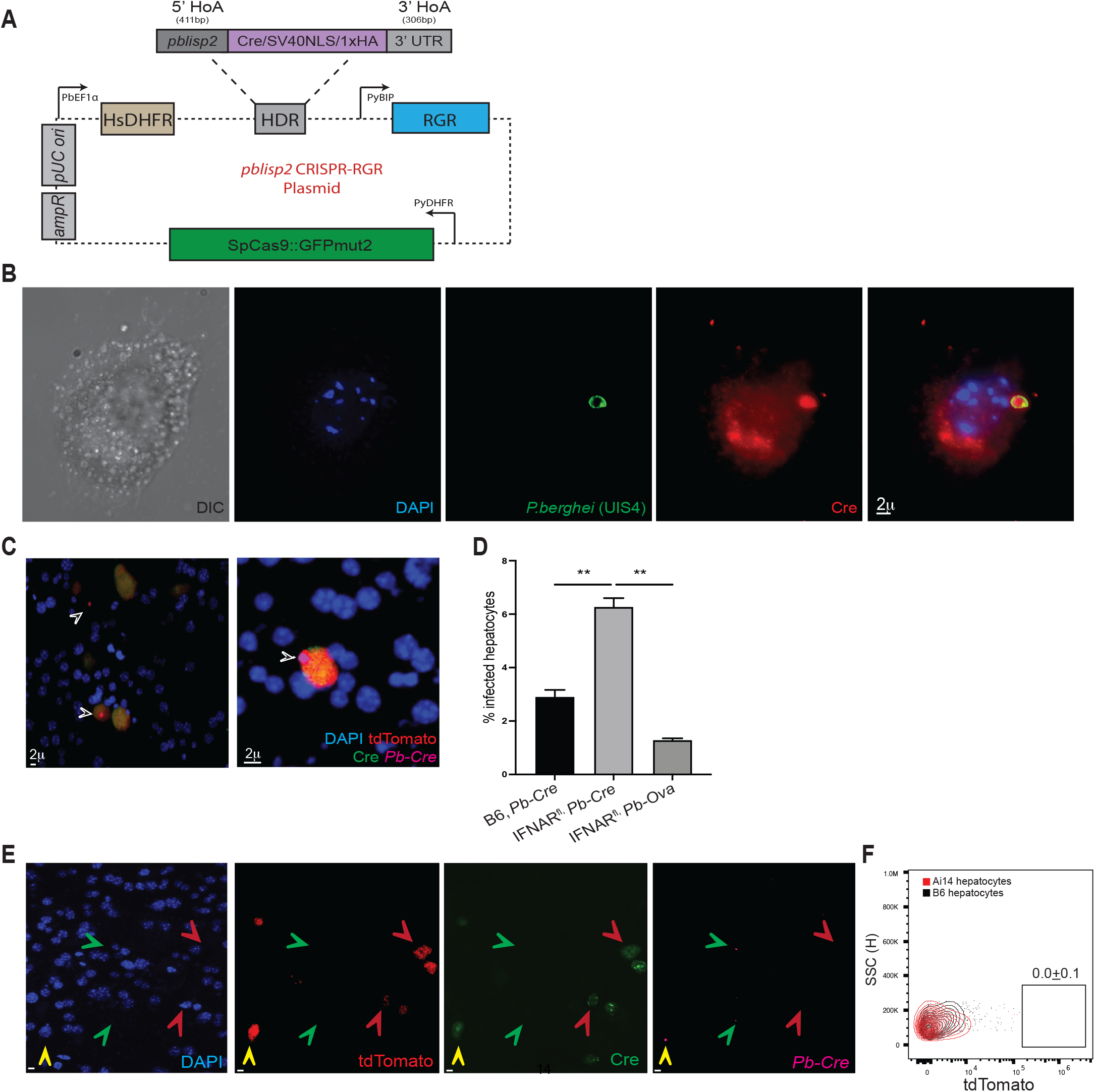
Cre recombinase transgenic *Plasmodium*. (A) Schematic of *pblisp2* CRISPR-RGR plasmid used to generate *Pb-Cre* parasites, indicating the key functional elements. A homology-directed repair (HDR) template was constructed that consisted of (1) 5’ homology arm (5’ HoA) comprising 411bp of the 3’ end of the *pblisp2* coding sequence, along with (2) coding sequence for a 2x(GGS) flexible linker, Cre recombinase, SV40 NLS, a single hemagglutinin tag (HA), and a stop codon, and (3) a 3’ homology arm (3’ HoA) comprising 306bp of the sequences immediately downstream of the *pblisp2* coding sequence. See the method section for more details. (B) Representative pseudo-colored deconvolved images indicating the expression and localization of Cre recombinase (anti-HA) in a *Pb-Cre* infected primary B6 mouse hepatocyte, 24h p.i. (C) Representative pseudo-colored confocal images indicating tdTomato expression, Cre localization in *Pb-Cre* (anti-Hep17) in primary Ai14 mouse hepatocyte culture. Arrows indicate infected hepatocytes. Image in the right panel shows a magnified view of an infected hepatocyte. (D) Frequencies of infected B6 or Ifnar1^flox^ primary hepatocytes infected with *Pb-Cre* or control *Pb-Ova* parasites when co-incubated with IFNα/β (100u/ml) for 36h. Data presented as mean + s.e.m and analyzed using ANOVA with Tukey’s correction. (E) Representative pseudo-colored confocal images indicating nuclear staining (DAPI), tdTomato expression, Cre localization or *Pb-Cre* (anti-Hep17) in primary Ai14 mouse hepatocytes. Green arrows indicate Pb-Cre infected hepatocytes, red arrow indicates tdTomato expressing hepatocytes and yellow arrow shows tdTomato expressing cells that have detectable *Pb-Cre* parasites in them. Related to Fig 2e, size bar = 2μM. (F) Frequency of tdTomato expressing hepatocytes in *Pb-Ova* infected B6 or Ai14 mice indicating baseline levels of tdTomato expression, at 24h p.i. Data presented as mean + s.e.m. All data represent ≥3 separate experiments. * P<0.05, ** P <0.01, n.s P≥0.05

## Materials and methods

### Mice and pathogens

C57BL/6 (B6), Ai14 mice, IfngKO, Alb-Cre and Ifnar1^flox^ mice were procured from Jackson Laboratories. cGAS KO and STING KO mice were provided by Dr. Rick Tarleton (University of Georgia). All mice were housed with appropriate biosafety containment at the animal care units at the University of Georgia. The animals were treated and handled in accordance with guidelines established by the UGA Institutional Animal Care and Use Committees. Luciferase expressing *P. berghei* (*Pb-Luc*) were obtained from UGA *SporoCore*. We generated *Pb-GFP, Pb-Cre* and *Pb-Ova* transgenic parasites. For infections, salivary glands of parasitized *A. stephensi* mosquitos were dissected and sporozoites (spz) were isolated as previously described(Kurup et al., 2019) and inoculated into cultures or injected intravenously (2×10^4^/mouse) in 200μL total volume. *P. berghei* parasites (ANKA strain) were used to generate *Pb-Cre and Pb-Ova* parasites. Transfection, selection, and cloning of *Pb-Cre* were performed as described in detail before(Janse et al., 2006; Kurup *et al*., 2019). In short, *in vitro* cultured schizont stages of *P. bergei* were transfected with the target plasmids using Amaxa parasite nucleofection kit II (Lonza), by employing the manufacturer’s protocol. The transgenic parasites were then inoculated into B6 mice and selected using pyrimethamine treatment. *A. stephensi* mosquitos infected with *P. berghei* ANKA, *Pb-Ova* and *Pb*-Cre were maintained at the University of Georgia insectary.

### *P. falciparum* gametocyte culture and mosquito infection

Gametocyte culture and mosquito infections were performed by modifying published protocols(Pathak et al., 2018). *P. falciparum* field isolate (CB132), was cultured with cryopreserved RBCs (Interstate Blood Bank, North Carolina, USA). Briefly, asexual and gametocyte cultures were maintained at 38°C with 5% O_2_ and 5% CO_2_ gas composition in an O_2_/CO_2_ incubator (Panasonic). The culture received daily media change with RPMI 1640 + L-Glutamine + 25mM Hepes + 50 ug/ml hypoxanthine (KD Medical, Maryland) and 5% heat-inactivated pooled human serum (Interstate Blood Bank, NC, USA). The asexual culture was kept between 0.5% to 5% parasitemia, and gametocyte culture was seeded at 0.5% parasitemia with cryopreserved O^+^ human RBCs (5% hematocrit). Gametocyte culture was assessed for exflagellation using a hemocytometer (INCYTO, South Korea) to count the number of exflagellation centers between 12-18 days post-seeding. Once deemed infectious during 12-18 days post-seeding, the cultures were offered to 150-250 3-7 days old female *Anopheles stephensi* mosquitoes as published before(Pathak *et al*., 2018). Infected mosquitoes were kept at 20°C ± 6 °C, 80% ± 5% relative humidity, and under a 12-hour day/night photoperiod schedule (Percival Scientific, IA, USA) with 5% dextrose (w/v) and 0.05% para-Aminobenzoic acid (w/v) (Thermo Fisher). Mosquito oocysts were checked at 10-days post-infection, and salivary glands were collected between 30-35 days post-infection.

### Generation of Pblisp2 CRISPR-RGR plasmid

A CRISPR-RGR plasmid for the insertion of Cre recombinase, SV40 NLS, and a 1xHA tag was created using pSL1394 as a base plasmid (Addgene # 129522). A homology-directed repair (HDR) template was constructed that consists of: 1) a 5’ homology arm (5’ HA) comprising 411bp of the 3’ end of the *pblisp2* coding sequence, 2) coding sequence for a 2x(GGS) flexible linker, Cre, SV40 NLS, a single HA tag, and a stop codon, and 3) a 3’ homology arm (3’ HA) comprising 306bp of the sequences immediately downstream of *pblisp2*. Additionally, a Ribozyme-Guide-Ribozyme sequence was synthesized as previously described(Walker and Lindner, 2019) that expressed two sgRNAs with perfect homology to the targeted region of *pblisp2*. Distinct and unique promoter and 3’UTR sequences were used for each transcriptional cassette to avoid recombination and removal of *Plasmodium* sequences in the plasmid.

### Murine primary hepatocyte culture, *in vitro* sporozoite infection

Primary hepatocytes were isolated from mice as described in detail before(Kurup *et al*., 2019). In short, in anesthetized (Ketamine/Xylazine (87.5/12.5 mg/Kg)) mice, the inferior vena cava was catheterized (BD auto guard, 22G) aseptically to perfuse the liver by draining the perfusate through the portal vein. A steady-state perfusion of the liver was performed first with PBS (4ml/min for 5 minutes), then Liver Perfusion buffer (4ml/min for 3 minutes, Gibco), and finally the Liver Digest Medium (4ml/minute for 5 minutes, Gibco). Digested liver was excised, single cell suspension made and resuspended in a wash solution of 10% v/v fetal calf serum (Sigma-Aldrich) in DMEM (Gibco). Hepatocyte fraction was recovered by centrifugation at 57g, from which debris and dead cells were removed by density gradient centrifugation using a 46% v/v Percoll (Sigma) gradient. Remaining cells were counted and resuspended in DMEM with 10% v/v FCS. Primary hepatocytes cultures were plated for 24h before infection. For luminescence measurement, 12×10^3^ cells were plated on collagen-coated 96-well plates in 50μl volume and incubated at 37°C, and 5% CO_2_, and then infected with 8 x10^3^ sporozoites per well. Cultures were further incubated for 24-36h to allow infection and liver-stage parasite development. Hepatocytes were stimulated with equal amounts of both murine IFNα and IFNβ (PBL) at varying concentrations, 1U=0.8pg.

### *P. falciparum* liver-stage Infection

Primary human hepatocyte infection in this study followed published methods(Roth et al., 2018). Briefly, cryopreserved primary human hepatocytes (BioIVT, NY, USA) were thawed two days prior to infection and seeded with 18,000 cells per well in a 384-well collagen-coated plate (Greiner, NC, USA). Hepatocyte cultures were maintained with daily media change with customized InVitroGro HI medium without dexamethasone (BioIVT, NY, USA) supplemented with 5% human serum (Interstate Blood Bank), 1:100 dilution of penicillin/ streptomycin/ neomycin antibiotic mixture (Gibco), and 1:1000 dilution of gentamicin (stock concentration:10 mg/mL, Gibco). The salivary glands of *P. falciparum* BD007 infected *A. stephensi* mosquitoes were collected between 18-20 days post-infection and were passed through a 30-gauge needle (Becton Dickinson). Total sporozoites were determined using a hemocytometer, and 24,000 sporozoites were inoculated into the hepatocyte culture. The plate was centrifugated at room temperature at 250 *xg* for 5 minutes with acceleration/break at 5. Hepatocyte cultures were kept at 38°C with 5% CO_2_ (Panasonic).

### Assessment of parasite burdens

Liver parasite burdens were assessed by quantitative real-time RT-PCR for parasite 18s rRNA in hepatocytes derived from mice challenged with *Plasmodium* sporozoites, as described before(Doll et al., 2016; Kurup *et al*., 2019). Total RNA was extracted from hepatocytes at the indicated time points post infection, using TRIzol treatment, followed by DNase digestion and cleanup with RNA Clean and Concentrator kit (Zymo Research). Two micrograms of liver RNA per sample was used for qRT-PCR analysis for *Plasmodium* 18S rRNA using TaqMan Fast Virus 1-Step Master Mix (Applied Biosystems). Data were normalized for input to the GAPDH control (hepatocytes) for each sample and are presented as ratios of *Plasmodium* 18s rRNA to host GAPDH RNA. The ratios depict relative parasite loads within an experiment and do not represent absolute values.

Blood stage parasitemia was assessed by flow cytometry as described before(Kurup et al., 2017). In the indicated time points, whole blood samples were stained with Hoechst 33342 (Sigma), anti-Ter 119 coupled to Phycoerythrin (PE) (Tonbo Biosciences) and anti-CD45 coupled to allophycocyanine (APC) (Tonbo Biosciences). Data were acquired on Cytoflex (Beckman Coulter) and analyzed using FlowJo X (Treestar).

The frequencies of *P. berghei*-infected hepatocytes *in vitro* were determined with CellTrace violet (CTV, Thermofisher) stained sporozoites and flow cytometry. Sporozoites were stained with CTV by modifying the manufacturer’s protocol, by incubating with 10mM CTV at 37°C for 20 minutes. The labeled sporozoites were then incubated for 5 minutes with 1mL FCS at 37°C to quench unbound CTV, washed once with media, and added to hepatocyte cultures. The frequencies of infected cells were determined by flow cytometry.

*Pb-Luc* infection loads were determined using Bright-Glo (Promega) as described before(Derbyshire et al., 2012). In short, in a 96-well plate, after 16h of infection cells were treated or not with the indicated equal concentrations of IFNα and IFNβ for additional 24h. Forty-four hours post infection, Bright-Glo reagent was added and parasitemia evaluated. The relative signal intensity of each plate was evaluated with Synergy H4 hybrid reader (Biotek) and Gen 5 2.0 System.

*P. falciparum* liver stage quantification was performed by immunofluorescence microscopy at 4 days post-infection. Infected hepatocyte treated with varying concentration of IFNα and IFNβ (1U=3.6pg) at 16h p.i.. At 4d p.i., cultures were fixed with 4% v/v paraformaldehyde (Alfa Aesar) for 10 min at room temperature and washed twice with PBS followed by permeabilization, blocking and staining with 0.03% v/v Triton-X (Acros), 1% w/v BSA (w/v) (Fisher Scientific) and 1:1000 dilution of mouse anti-GAPDH 13.3 (stock concentration 1mg/ml, European Malaria Reagent Repository, UK) overnight at 4 °C. A secondary antibody mixture was made using the same permeabilization/blocking solution with 1:1000 dilution of anti-mouse Alexa Flour 488 (Thermo Fisher) overnight at 4 °C. The wells were then stained with 10 μg/ml Hoechst under room temperature for 1 hr. Fixed and stained samples were stored in PBS for quantification. The number of parasites in each well was imaged using ImageXpress and quantified using MetaXpress software (Molecular Devices, CA, USA). Images were acquired using FITC, and DAPI channels at 10X magnification, resulting in each well of a 384-well plate offering 9 fields of view, tiled together. After image acquisitions, hepatocyte nuclei were counted with the DAPI channel. Parasites were counted with the FITC channel and were identified by area, mean intensity, and cell roundness.

### Therapeutic regimens

The following *in vivo* treatment regimens were used in this study: DMXAA (5,6-dimethylxanthenone-4-acetic acid, Invivogen): 20mg/Kg intraperitoneal (i.p.), 1d p.i. Poly I:C (Invivogen): 100ug/mouse, hydrodynamic i.v., 1dpi. Murine IFNα and IFNβ: 3.3μg/Kg i.p. 12h p.i. and 24h p.i.

### Microscopy

*Plasmodium*-infected hepatocytes cultured (30h p.i.) on collagen coated cover-slips were fixed with 4% paraformaldehyde in PBS for 10 minutes. The samples were washed with PBS and coincubated with 0.25% v/v Triton X-100/ PBS (Fisher Bioscience) for 10 min at room-temperature to permeabilize plasma membranes. Samples were subsequently blocked with 1% w/v BSA/PBS (60 min), stained with specific antibodies that delineate the PVM (anti-UIS4 or Hep17, in house generated), anti-IRF-9 (Clone 6F1-H5, Millipore Sigma), anti-HA (Clone 16B12, Biolegend) anti-Cre Recombinase (Clone poly9080, Biolegend) or 4’,6-Diamidino-2-Phenylindole, Dihydrochloride (DAPI, Sigma Aldrich) for 60 mins. The samples were washed thrice with PBS, probed with fluorophore conjugated secondary antibodies, washed thrice with PBS again and visualized in Applied Precision DeltaVision Microscope System I (Olympus IX-71) and processed using SoftWorx software.

### Transcriptome analysis

*Pb-GFP* infected or naïve (from uninfected cultures) hepatocytes derived from cultured prepared as described above were FACS sorted and re-suspended in TRIzol. RNA was purified with RNAeasy kit (Qiagen) as described previously(Vijay et al., 2017). RNA was assessed for purity and quality using an Agilent 2100 Bioanalyzer and transcriptome analysis performed on Affymetrix GeneChip Mouse Transcriptome array 1.0. Data output was analyzed and visualized using Affymetrix Expression Console v1.4.4.46, Applied Biosystems Transcription Analysis Console v4.0.1.36 and Interferome v2.01 (www.interferome.org) and submitted to NCBI Gene Expression Omnibus.

### Statistical analyses

Statistical differences between two study groups were evaluated using two-tailed t-tests. Statistical differences between more than two study groups were evaluated using a one-way ANOVA with the indicated corrections applied. Statistical significance was calculated based on the numbers of biological replicates as indicated in the figures or figure legends and assigned as *p < 0.05, **p < 0.01, n.s: p>0.05. Statistical analyses were performed using Prism 9 software (GraphPad).

## Author contributions

C M-d-S, KP, MW, JS, RV, SEL, and SPK designed, performed or analyzed the experiments. SEL and DEK provided vital resources. C M-d-S and SPK wrote the manuscript. The study was supported by NIH R21AI130692 to SEL, NIH R21AI144591 to DEK, T32AI1060546 to JS and UGA research startup to SPK.

## Acknowledgements

We want to acknowledge the contributions of Dr. John Harty (University of Iowa) for offering vital insights pertinent to the study, and along with Dr. Rick Tarleton (University of Georgia) for critiquing the manuscript. We acknowledge the contributions of UGA Office of animal research, UGA CTEGD central microscopy core, UGA CTEGD central flow core and Iowa Institute of Human Genetics for providing valuable access to their facilities, as well as the technical support provided by Drs Diego Huet and Ronald Etheridge (UGA)

